# Assessing the metabolomics “dark matter” by a detectable khipu model

**DOI:** 10.1101/2025.02.04.636472

**Authors:** Yuanye Chi, Joshua M. Mitchell, Shujian Zheng, Maheshwor Thapa, Shuzhao Li

## Abstract

There is no consensus on how to interpret the large number of unknown features in untargeted metabolomics, which are sometimes referred as the “dark matter”. Are these features real compounds or artifacts? Understanding this problem is critical to the annotation and interpretation of metabolomics data and future development of the field. We propose a “detectable khipu” model here, to show that compounds exhibit ion group patterns that depend on their abundance. We apply this model to a systematic analysis of 61 representative public datasets from blood LC-MS metabolomics, the most common data type in biomedical studies. The results indicate that majority of abundant features have identifiable ion patterns, and in-source fragments contribute to less than 10% of features. Each dataset detects 1∼2,000 high confidence compounds, over half of which are unknown. The major knowledge gap in LC-MS metabolomics is therefore not the methods of grouping ions or counting fragments, but the identification of unknown compounds.

---

Metabolomics aims to measure all small molecules in biological systems. Currently, the majority of metabolomics data in biomedical research are produced using LC-MS (liquid chromatography coupled mass spectrometry). In over a decade, no consensus has been reached on how to interpret the unidentified features in untargeted metabolomics, which are typically majority in numbers and sometimes referred as the “dark matter” (daSilva 2015; Monge 2019). It is important to understand this knowledge gap. If these unidentified features are analytical artifacts, we should not waste resources to further characterize them; if they are real compounds, it is mandatory to establish their biological relevance. This problem underlies the analytical coverage of small molecules in biomedical research (Kind 2009; Uppal 2016), approaches to metabolite annotation (Domingo 2018; Chaleckis 2019; Metz 2025), mapping reaction pathways (Zamboni 2015; Artyukhin 2018) and the promise of applying metabolomics and exposomics to precision medicine (Wishart 2016; David 2021; Banbury 2025). As the stakeholders of metabolomics range across many disciplines, including chemistry, biology, informatics, biotechnology and medicine, data-based evidence is critical for decision making.

Multiple factors contribute to this metabolomics “dark matter”. Modern mass spectrometry (MS) brings tremendous new power into chemical analysis, enabling the detection of compounds that were previously elusive due to limited separation, abundance or stability (Xian 2012; Uppal 2016; Lai 2024). Biology has knowledge gaps and new biochemical mechanisms continue to be discovered. E.g. hundreds of unknown metabolites can be affected by one enzyme (Artyukhin2018; Heremans2022). Such results suggest that characteristic metabolite modifications may be as widespread as protein posttranslational modifications. Our cohabitation with the microbiome results in sharing many metabolites from under-characterized microbial species (Peisl2018). Challenges exist in data acquisition, processing and interpretation.

Contamination is common in tandem mass spectra (MS/MS) from biological samples (Stancliffe 2021). Artifacts can be generated by data processing software when sensitivity is not matched to detection confidence (Li2023b). Because isotopologues, adducts and fragments are commonly measured in the MS data, it is an apt question if the measured metabolome is inflated by degenerate features (Mahieu2017; Wang2019; Li2023a). In-source fragments (ISFs), generated when molecules are broken down during the process of ionization, became a focus of recent debates. Giera et al (2024) reported that ISFs accounted for over 70% of MS/MS features in METLIN, one of the leading spectral databases, suggesting that ISFs could be a significant portion of the “dark matter”. El Abiead et al (2025) calculated the presence of ISFs using a different compound library, reporting less percentage but still two fragments on average per protonated compound. However, both studies focused on data from chemical standards, which are primarily used for metabolite annotation, different from complex biological samples.

Here, we present a “detectable khipu” model to explain how many compounds are present in untargeted LC-MS metabolomics. Systematic analysis of 61 representative datasets from blood LC-MS metabolomics indicates that the contribution of ISFs is small, but the majority of abundant features have identifiable ion group patterns. Each dataset detects 1∼2,000 high confidence compounds (defined by valid ^13^C/^12^C patterns), over half of which are unknown. The major knowledge gap is therefore not the methods of grouping ions or counting fragments, but the identification of unknown compounds.

## Results

### A “Detectable Khipu” model of mass spectrometry data

High-resolution mass spectrometers are now routinely employed in metabolomics. The data often contain multiple ions per compound, in the forms of isotopologues, adducts and fragments (conjugates or multimers are possible but rare). As a result, the number of features is not a direct account of number of compounds, thus a source of longstanding debates on the real number of detected compounds (Mahieu2017; Wang2019; Uppal2016). Pre-annotation, the process of grouping ions into tentative compounds, depends on signature mass differences, e.g., 1.0034 between ^13^C (carbon 13 isotope) and ^12^C and 21.9819 between Na^+^ and H^+^ adducts (**Figure 1a**). Many software tools perform some variation of pre-annotation (Kuhl 2012, Mahieu 2016, DeFelice 2017, Uppal 2017, Kachman 2020, Chen 2021). The khipu algorithm was introduced to perform de novo, global pre-annotation using a generalized data structure (Li2023a). In short, a khipu aligns coeluting ions on a grid of isotopologues and modifications, where modifications include adducts and fragments (**Figure 1a, Figure S1**). The khipu construction leads to an inferred neutral mass, and the data structure enables downstream computing.

**Figure 1.**
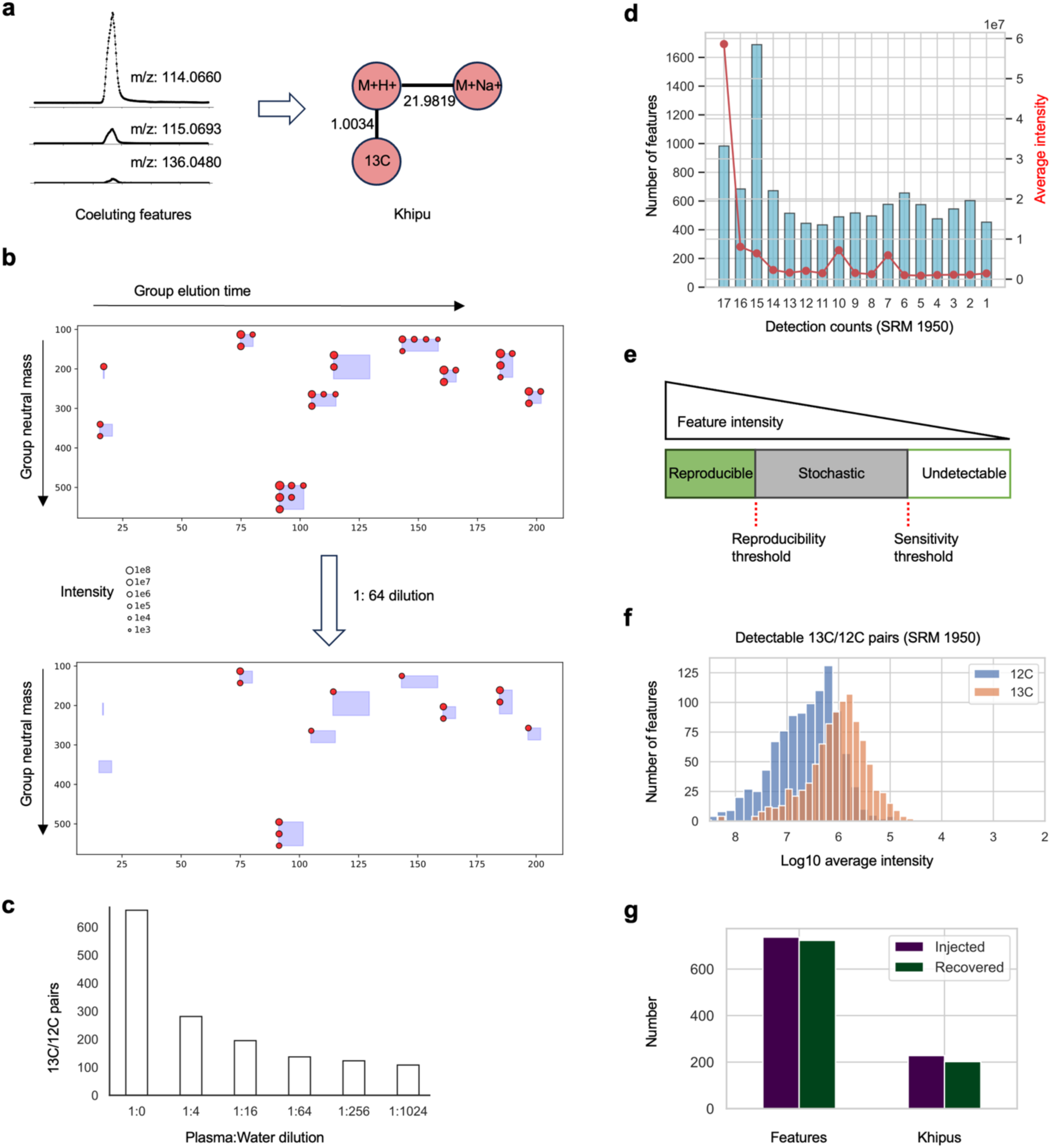
A detectable khipu model for feature groups in untargeted metabolomics. **a.** A khipu is a group of features that belong to the same tentative compound, represented as a tree of isotopes and modifications. **b.** The detection of khipus as a function of concentration. Each shaded rectangle marks a khipu in the top panel, and each red dot a feature. After dilution, many features are not detectable and their khipus are not identified (bottom). **c.** Number of 13C/12C feature pairs as a function of dilution factor. **d.** Detection frequence of LC-MS features in a dataset of 17 repeated injections of NIST SRM 1950 reference material. Each bar represents a bin of features. The y-axis on the right is average signal intensity of each bin of features (red). **e.** Dependency of feature detection and quality on feature intensity. **f.** Distribution of intensity of 13C/12C feature pairs in the SRM 1950 dataset. **g.** Khipu performance tested on a synthetic dataset, where 737 features from 228 compounds were injected into a biological dataset. Using parameters in this study, 723 features of 202 empirical compounds were correctly recovered by the khipu algorithm.

The presence of a khipu indicates high-confidence detection of a compound (not necessarily of biological origin). In theory, any compound can have a khipu of multiple ions. However, ions under a certain abundance level are not detected in experimental data. To illustrate this concept of “detectable khipus”, we performed untargeted LC-MS metabolomics on a serial dilution of human plasma by water. Many ions as part of khipus are no longer detected after 1:64 dilution, and many khipus become singletons (**Figure 1b**). The number of detected ^13^C/^12^C feature pairs decreases with the dilution factor (**Figure 1c**). This suggests that molecular abundance is a major determinant of finding the khipu patterns, and singletons are incomplete data of khipus due to experimental detection limit. An alternative explanation of singletons is the failure of computational methods to account for a connection to other detected ions, which is further investigated in this paper.

The “detectable khipus” apply to both a single mass spectrum (commonly used in MS/MS) and LC-MS features (**Figure S1**). It is important to note that features of interest in a spectrum are mass peaks (**Figure S1a**), while in the current practice of LC-MS, features are elution peaks that are observed in consecutive spectra (**Figure 1a, Figure S1b**). The latter filters out many spurious data points.

### The long tail of data quality in untargeted metabolomics

Not all features in untargeted metabolomics are reproducible. An experiment is typically optimized for a subset of metabolites, while including low-quality measurements of many others. We composed a dataset using 17 repeated measurements of a common reference sample (NIST SRM 1950), each from a separate batch, and plot here the detection frequency of each LC-MS feature (**Figure 1d**). Only 983 out of all 10811 features, about 9%, are present in all 17 injections thus considered reproducible, while the remaining features are detected in varying subsets. The reproducible features are of much higher signal intensity than the others (**Figure 1d, Figure S2a**). Other factors, such as metabolite stability, contribute to reproducibility, but a practical schema can be postulated that reproducible features are at the top end of intensities; many low-intensity features are not reproducible; and the experimental detection limit cuts off low-abundance ions (**Figure 1e**). In this dataset, most reproducible features are over 1e6 in intensity (**Figure S2a**), similarly indicated by plotting ^13^C/^12^C paired features (**Figure 1f**). On similar platforms, these high-intensity features account for less than 10% of all features in untargeted LC-MS data (**Figure S2b, c**), suggesting a long tail of low-quality features in every dataset. The feature quality can be quantified by signal-to-noise ratio (SNR) and peak shape (goodness of fitting to a Gaussian curve), as applied below, but they tend to correlate with intensity levels.

In the above experiment, the feature space is defined by one sample. However, heterogenous biology and chemical exposures are present in human population studies, leading to features that are expected only in subset of samples. These low-frequency features therefore increase the numbers of total features as well as low-quality features.

### Evaluation of khipu performance using a synthetic dataset

The performance of the khipu algorithm depends on the list of search patterns, as well as data complexity, how the mass values are resolved and elution peaks are separated. The latter can vary significantly between experiments and laboratories. To obtain a baseline metric on its performance, we have created a synthetic dataset to test pre-annotation. In a library of 228 authentic compounds based on 50 LC-MS acquisitions, 737 features were identified as ions from these compounds. These features were computationally “injected” into a biological dataset of the same LC-MS method (Methods) to serve as ground truth (Suppl File S1). As shown in **Figure 1g**, 723 of the 737 features were recovered by the khipu algorithm, and 202 of the identified khipu feature groups match to the authentic compound groups. This indicates that the method performs reasonably well in complex biological datasets.

### Systematic survey of 61 public datasets

The most common platforms of LC-MS employ orbital mass analyzers (Orbitrap) or time-of-flight (TOF) analyzers, using positive or negative electrospray ionization (ESI). Even with similar methods, the data often have considerable variations cross labs. To survey the field, we retrieved 61 LC-MS metabolomics datasets from public repositories, including 45 Orbitrap and 16 TOF datasets (**Figure 2a, Table S1**). For consistence of data analysis, only studies of human plasma or serum samples are included, and large datasets are down selected to about 100 samples, because low-frequency features accumulate by increasing sample number. We use the asari software (Li2023b) for processing raw data to features of unique m/z (mass-to-charge ratio) and retention time (RT). Asari is designed with explicit data models that report all features above a quality threshold (SNR of 2, peak shape of 0.5). The median number of reported features in the Orbitrap studies is around 50,000, of which about a quarter to a third have SNR > 5 and peak shape > 0.9. The numbers of ^13^C/^12^C pairs are typically between 1,000 to 3,000 in these metabolomic datasets (**Figure 2b**). The presence of a ^13^C/^12^C pair indicates high-confidence detection of a khipu; additional isotopes and adducts do not change the total count of khipus or estimated number of compounds (**Figure 1b**). But fragments can have their own isotopes and adducts, thus inflate the count (**Figure S1**). Therefore, in-source fragments are investigated below using two approaches: observed mass differences in LC-MS and MS patterns that match to MS/MS data.

**Figure 2.**
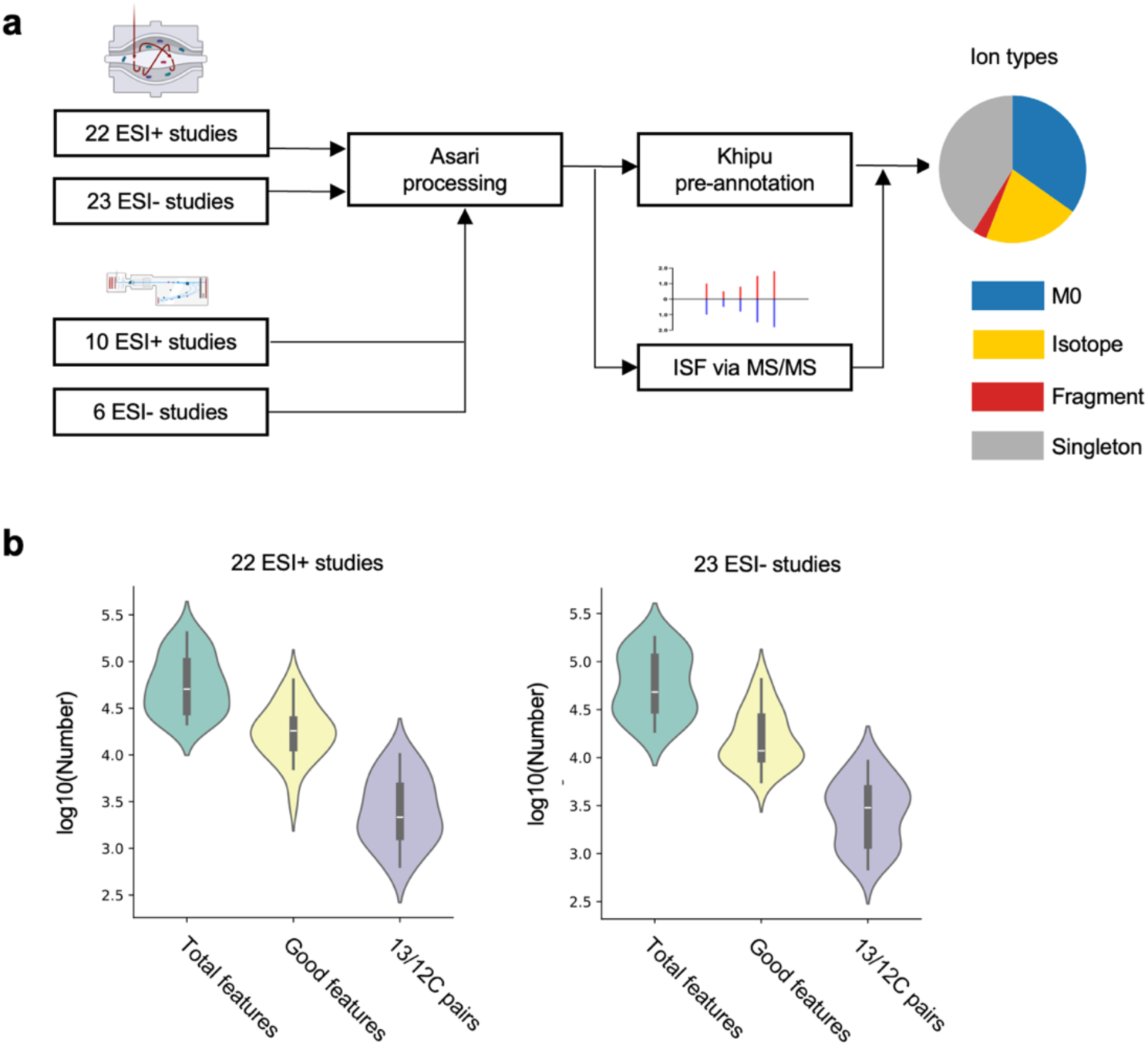
Processing and analysis of 61 LC-MS studies **a.** Schematic of pre-annotation analysis, from orbital or time-of-flight instruments. ESI: electron spray ionization; ISF: in-source fragment. M0: the isotopic form of ^12^C only. The results from 16 TOF datasets are shown in supplements. **b.** Across the 22 Orbitrap studies using positive ionization, the median number of features is 50,771, the median number of good features 18,117, and the number of matched ^13^C/^12^C pairs 2,176. Similar numbers are observed across the 23 negative ionization Orbitrap studies with 48,212 features, 11,783 good features, and 3,008 ^13^C/^12^C pairs. Good features are defined as having SNR > 5 and peak shape > 0.9 when fitting to a gaussian curve.

### Common mass differences in LC-MS and application to ISF estimation

The pre-annotation approach depends on the list of mass differences underlying the ion patterns. The khipu software includes default lists of isotopes and adducts based on prior data. To be comprehensive, we also systematically calculated all common mass differences between coeluting features in these current LC-MS datasets (**Table S2-5**). The coelution window here is defined as two standard deviation of RT differences of all ^13^C/^12^C pairs. Not surprisingly, the m/z difference between ^13^C/^12^C isotopologues is by far the most frequent observation in both ESI+ and ESI- data (**Figures 3a, S3a**). The most frequent mass differences, excluding known isotopic and adduct patterns, can be considered as candidates for ISFs (also referred as neutral losses).

**Figure 3:**
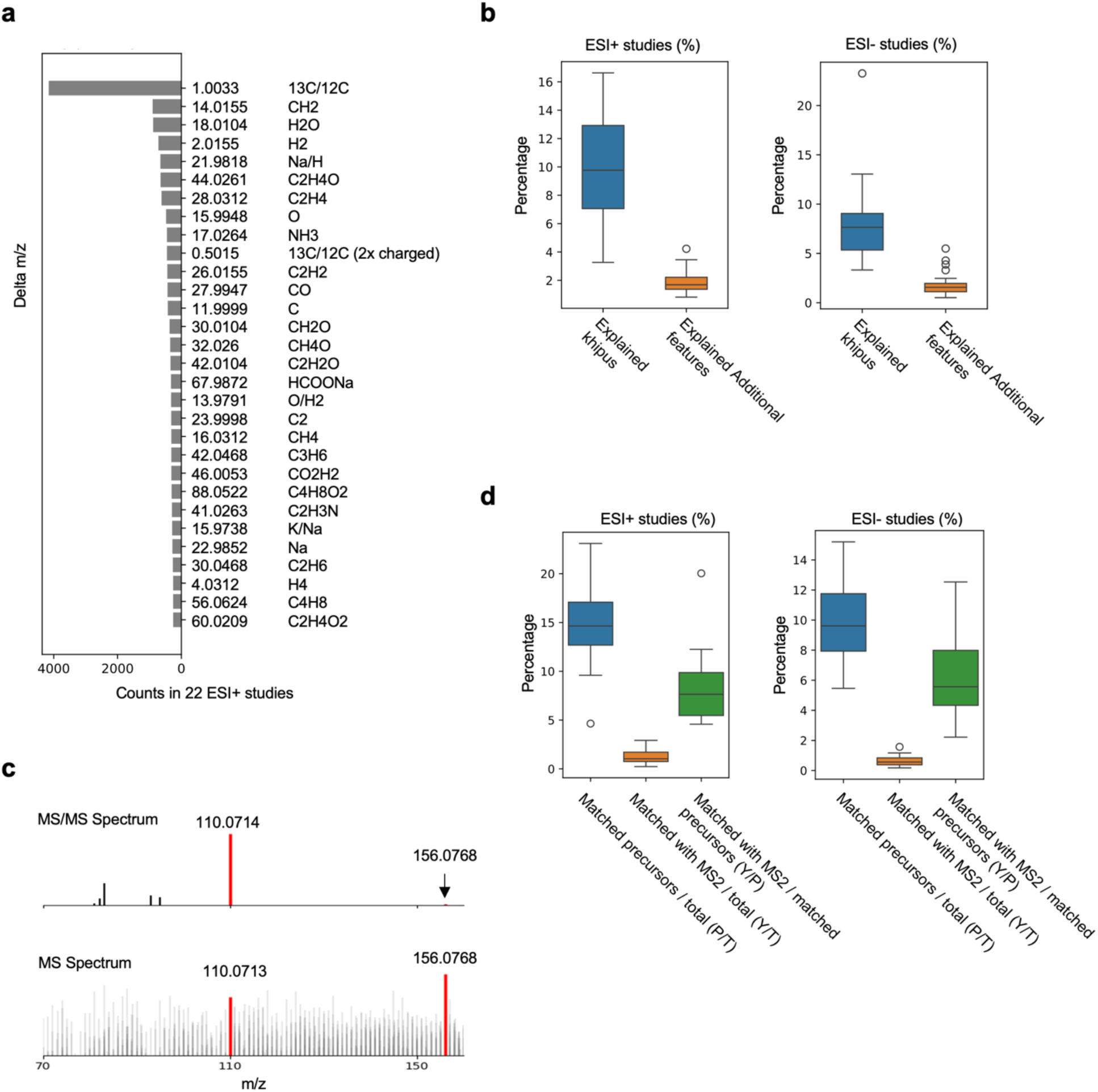
In source fragments account for less than 10% of LC-MS features. **a.** Most frequence delta m/z values in positive ionization Orbitrap datasets. **b.** The percentages of khipus and additional features explained by candidate fragments, in Orbitrap positive and negative ionization datasets, respectively. **c.** Example of matching MS/MS and MS spectra for searching ISFs. Top: MS/MS from MoNA. Bottom: sample CHCL3_20min_B from dataset MTBLS1465_HILICpos ppm5_3505731, MS scan number 1408. **d.** The percentages of features matched to MoNA MS/MS spectra across the 45 LC-MS studies, using positive and negative ionization respectively. In each boxplot, 1^st^ column is the percentage of features matching a precursor m/z value (P) over total features (T), the 2^nd^ column the percentage of features matching at least one MS/MS fragment (Y) over total features (T), and the 3^rd^ column as Y over P.

To assess the impact of ISFs on full metabolomic profiles, we compare the pre-annotations without and with consideration of ISFs. Without ISFs, the khipu algorithm is applied to the LC-MS datasets using a conservative set of isotopologues and adducts (**Methods**), to produce a set of khipus and singletons per dataset. The numbers of khipus and singletons can be impacted by ISFs, but ISFs do not impact relationships between isotopologues and adducts. Next, to consider ISFs, the candidate ISFs (**Table S6, Figures 3a, S3a**) are searched between coeluting khipus, and between coeluting singletons and khipus (**Figure S3b**). The comparison between two steps reveals that in the ESI+ data, about 500 khipus and 700 singletons can be explained by the candidate ISFs (**Figure S3c**), which correspond to about 8% of khipus and 1% of all features, respectively (**Figure 3b**). In the ESI- data, the candidate ISFs explain about 5% of khipus and 1% of extra features (**Figure 3b**). While these results do not rule out additional ISFs among singleton features, singletons are of lower intensity and expected to contain even less percentage of detectable fragments.

### MS/MS based search corroborates less than 10% ISFs in LC-MS data

Typically, ESI is used in MS to preserve intact molecules by minimizing fragmentation. In contrast, MS/MS techniques fragment molecules intentionally by a collision energy, and the fragments serve as a fingerprint of the molecule. If ISFs of a molecule are present in LC-MS data, they must a) have the same retention time as the intact molecules, and b) have m/z values that are similar to those in the corresponding MS/MS spectra. In Giera et al (2024), MS/MS data at zero collision energy were used to mimic MS data. We take a similar strategy here to compare an MS/MS database (MoNA2024) to the LC-MS metabolomics data, requiring both the precursor and at least one MS/MS fragment to be observed in MS data (**Figure 3c**). Even though the MoNA database does not fully cover the LC-MS metabolomics data, the number of matched MS/MS fragments relative to the matched precursor ions should indicate the prevalence of ISFs. We parsed out 13,672 compounds from MoNA MS/MS spectra. They typically match to 5∼10,000 features by precursor m/z values (**Figure S3d**). Of these matched precursor ions, about 7% have at least one matched MS/MS fragment in the elution window for positive ESI data, about 5% for negative ESI (**Figure 3d**). These results confirm the above khipu analysis that ISFs impact less than 10% of LC-MS features. The results are expected to include false positives (thus overestimation), as a) MS/MS fragments generated by higher collision energies are not expected to be in the LC-MS data; and b) matched features are not necessarily generated by ISF.

### Most features in untargeted metabolomics conform to the “Detectable Khipu” model

As indicated in **Figure 1**, khipu patterns are detectable in high-abundance features but high-abundance features are a small fraction of all data. To test the dependency of pre-annotation on feature abundance, we rank the features in each dataset by intensity and calculate the khipu explained features in serial bins of 1000 features. For the top 1000 features in Orbitrap ESI+ data, the median number of khipu-explained features is 856, which decreases in lower-intensity bins (**Figure 4**). Similar result is seen in Orbitrap ESI- data (not shown). The same trend is observed when all features are analyzed by intensity quartiles (**Figure S4a, Tables S1, S7**).

**Figure 4:**
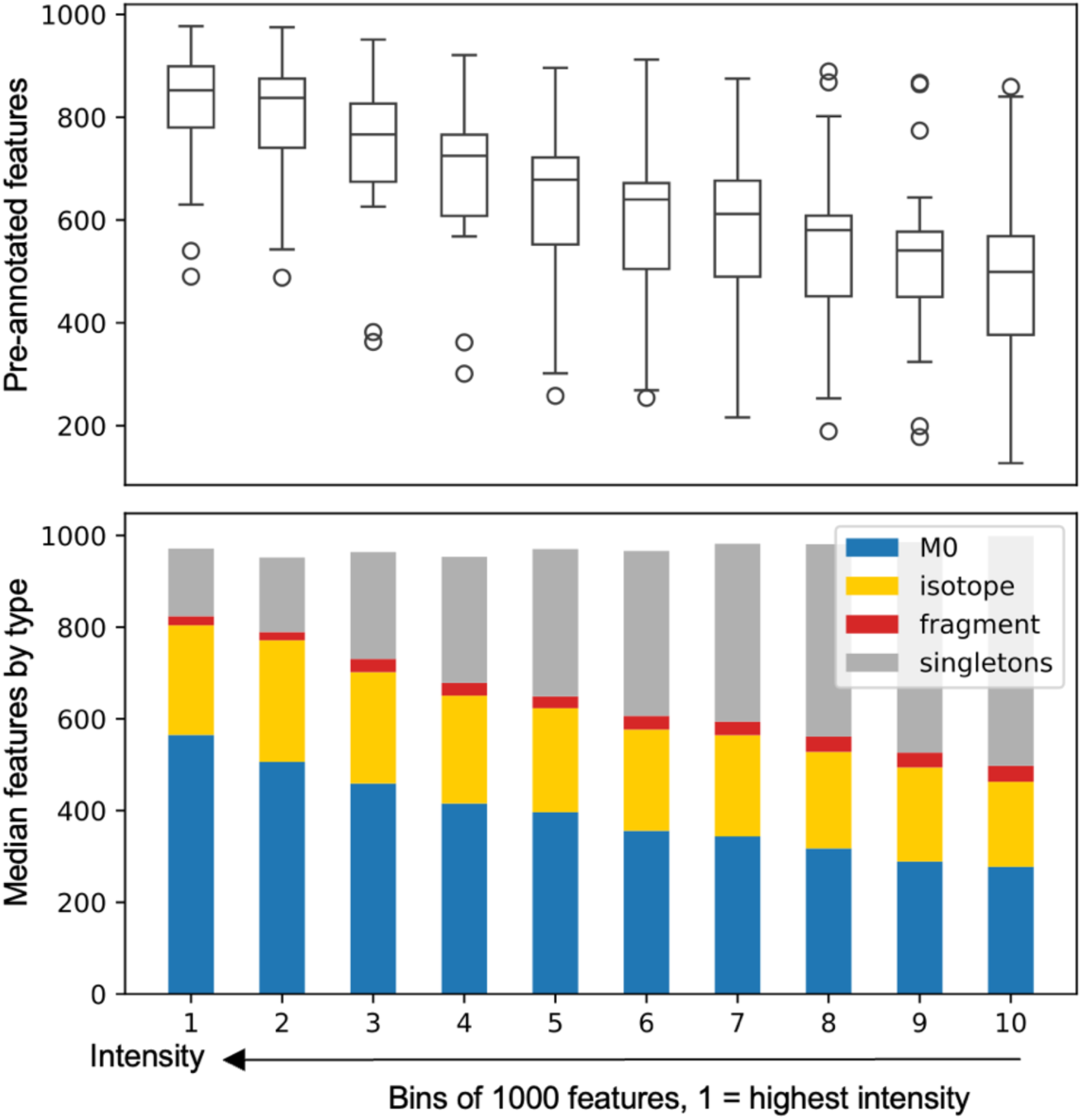
Features explained by khipus in each intensity bin in 22 Orbitrap ESI+ studies. Each feature table is sorted by feature intensity and the top 10 bins are shown here. Each bin represents 1000 features. Top: total number of khipu explained features. Bottom: median values per ion type. All adducts of an isotope other than M0 are considered as isotopes.

Within the top quartile, the singletons, i.e., features not explained by khipus, have significantly lower intensity than monoisotopic (M0) features (**Figure S4b**). To test the impact of peak alignment on the results, we selected one random sample (thus avoiding peak alignment) from each dataset to repeat the full data processing and pre-annotation, which returned similar results (not shown). Within the top bin in **Figure 4**, over 80% features are explained by khipu in most studies, while the exact number correlates more with high-quality features than with total feature number (**Figure S4c**). Taken together, the most high-abundance features in the surveyed 61 studies are explainable by khipu pre-annotation, while the full feature tables conform to the “detectable khipu” model, with increased fraction of singletons in lower intensity features.

### Incomplete compound annotation in blood metabolomics

How many compounds are measured in these plasma/serum metabolomics datasets? How many are known? In the public Orbitrap datasets, each contains typically 5∼6,000 unique khipus, averaging three features per khipu. The median number of high-confidence khipus (with valid ^13^C/^12^C patterns) is 1,338 for the ESI+ datasets, and 2,028 for the ESI- datasets (**Figure 5a**). Less than half of the high-confidence khipus have matches to the HMDB, the leading metabolite database (Wishart2022), by neutral mass (**Figure 5a, b**). Because many false positive matches are expected by searching neutral mass and HMDB is not tissue specific, the real compound annotation is much less.

**Figure 5:**
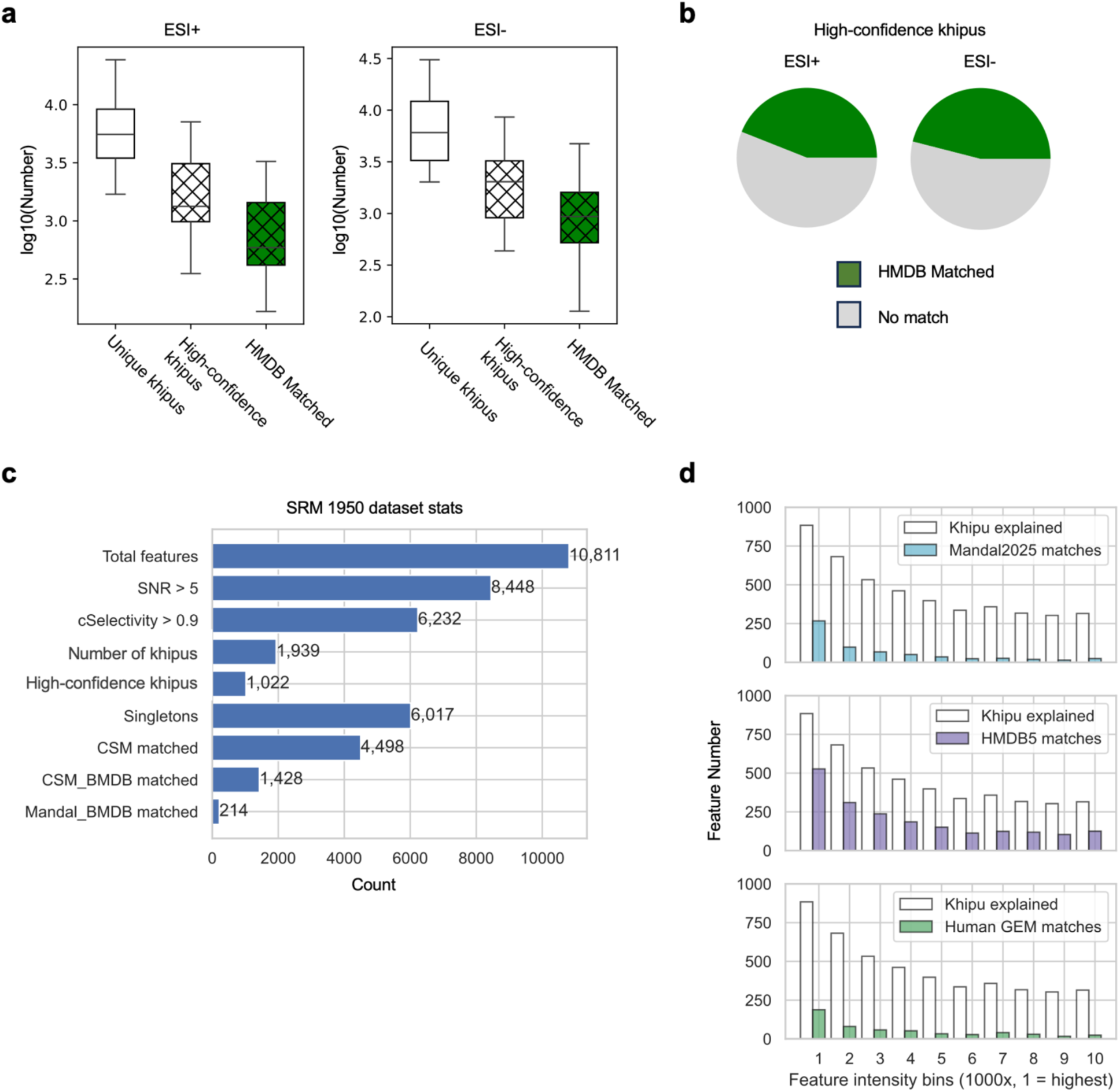
Annotation coverage in untargeted LC-MS metabolomics. **a.** Numbers of unique khipus, high-confidence khipus (those of valid 13C/12C patterns) and HMDB matched (by neutral mass) high-confidence khipus in Orbitrap positive and negative ionization datasets, respectively. **b.** Fractions of high-confidence khipus with HMDB matches. **c.** Summary statistics on the SRM 1950 dataset (same data as in Figure 1d). Here, cSelectivity is a measure of clean chromatography (Li2023b). CSM is the consensus serum metabolome (Chi 2025); BMDB is the annotation database in CSM, which overlaps with the Mandal et al (2025) result (Mandal_BMDB contains 706 compounds). The CSM annotation method returns 214 compounds matched to the list by Mandal et al. **d.** Comparison of khipu explained SRM 1950 features to numbers of features matched to annotation sources. From top to bottom: Mandal et al (2025) as known compounds in the SRM 1950 material, HMDB and a human genome scale metabolic model (Robinson2020).

The question of annotation coverage can be addressed on our SRM 1950 data more specifically. Our SRM 1950 data contain 10,811 features by a HILIC ESI+ method, of which 1,022 are high-confidence khipus (**Figure 5c**). Compared to a recent large-scale construction of a human consensus serum metabolome (CSM, Chi 2025), 4,498 features are also observed in the CSM, i.e. in other public datasets (**Figure 5c**). By examining serial bins of 1000 features by abundance ranks in this dataset (using the same method as in Figure 4), 884 features in the top bin can be explained by khipu pre-annotation (**Figure 5d**). A partial ground truth on the SRM 1950 reference material was recently provided by Mandal et al (2025), listing 922 HMDB compatible compounds by combining multiple analytical methods. By neutral mass comparisons, we match the khipus in SRM 1950 to three different sources of annotation: Mandal et al, HMDB and a human genome scale metabolic model (Robinson2020). Of the 884 khipu explained features in the top bin, 267 are matched to Mandal et al; the number decreases greatly for the lower-intensity bins (**Figure 5d**). Overall, the numbers matched to genome scale metabolic model are smaller, while those to HMDB are greater (**Figure 5d**). Since all matches are based on mass, these numbers are not precise but provide an upper limit of annotation.

These results indicate that the majority of compounds in LC-MS blood metabolomics are unknown. To examine the impact of analytical contaminants, 996 features from the blank control samples are removed from the feature table, which leads to the reduction of about 10% of khipus (**Figure S5** compared to Figure 5c). The blank filtering casts little impact on named compound annotation, reducing only 35 out of 1428 in CSM and 1 out of 214 in Mandal et al.

## Discussion

This study provides a systematic survey of blood LC-MS metabolomics data using the “detectable khipu” model. High-quality features are a small subset of all features, and the number of compounds is much smaller than the number of features. In all datasets, the most abundant features are largely explainable by khipu pre-annotation. This suggests that the interpretation of less abundant features is just limited by concentration and ionization efficiency. Therefore, this “detectable khipu” analysis offers a plausible explanation of the “dark matter” of metabolomics. The corollary is that the overwhelming majority of LC-MS metabolomic features from human blood samples are part of detected khipus (high abundance) or undetected khipus (low abundance), thus likely to be from real compounds. Most of these compounds are not matched in HMDB and considered unknown. We do not assume all the compounds are of biological origin. A compound should only be considered as a biological metabolite based on evidence of its activity. Contaminants from sample processing and instrument analysis can contribute to the detected compounds. Our result showed about 10% khipus from contaminants in the SRM dataset. However, the percentage would vary by sample size and many other factors. It is not feasible to track blank controls in most public datasets, but the CSM analysis (Chi 2025) indicated that contaminants are not common cross datasets, presumably because they vary between laboratories.

The khipu approach, like other computational tools for pre-annotation, relies on a catalog of mass differences. A new catalog of mass differences was produced by this study from a large collection of public datasets. However, the catalog is far from exhaustive. Based on mass differences, the method cannot distinguish fragments of constant sizes and can confuse in-source reactions with biological reactions. While we demonstrated that khipu successfully recovered spike-in signals in the synthetic data, the performance will vary cross experiments, dependent on chromatographic separation and mass resolving power. The approach explained 80% not 100% of the top features, suggesting room for improvement, such as more comprehensive list of m/z delta values and considering conjugates. Edge cases also exist, e.g. some isotopic features may be out of scan range; misalignment in retention time leads to pre-annotation errors. More structure-specific modeling in the future can also improve the approach.

While previous studies examined the numbers of compounds and fragments in MS/MS data or in isolated experiments, this work surveys a sizable data collection cross many labs, and examines MS1 profiles directly by two orthogonal approaches. The results lead to realistic numbers of reliable measurement of compounds in untargeted metabolomics. Some researchers are already aware of this, but the data-driven evidence presented here helps clarify it for the broader community. Likewise, the illustration of the long tail of low-quality data highlights the need for caution when pursuing analytical coverage, sensitivity, and reproducibility. The proposal that singletons are incomplete khipus is a conceptual advance. It suggests that singletons in one study may be observed as khipus in another study under more favorable analytical conditions. This framework shall facilitate cumulative annotation at the community level.

Our results indicate that less than 10% of features in blood LC-MS data are explained by ISFs, while the number may vary between studies and for different compound classes. This is consistent with previous LC-MS (Guo2021) and LC-MS/MS (He2020) studies, but quite different from the numbers reported by Giera et al (2024) and El Abiead et al (2025). However, results from all studies have a consistent explanation. ESI can generate many fragments but of low intensity (also pointed out by El Abiead et al, 2025). In the analysis of authentic chemical standards, many ISFs are detected by mass spectrometer because there is little background and the input concentration is high, while the percentage varies by platforms. There are often over 1 million mass peaks per sample in LC-MS metabolomics, where data are processed into features based on elution peaks. Most ISFs are not observed in LC-MS feature detection because a) signals inconsistent cross scans are filtered out by elution peaks, and b) low intensity ISFs are outcompeted by more abundant molecular species in the complex matrix of biological samples.

Several precautions may protect us from confusions in the field. Metabolomics moves the traditional analytical chemistry to the-omics scale of complex data. Statistics in spectral databases is difficult to extrapolate directly into LC-MS metabolomics on biological samples. Explicit data models are important to define the purpose and context of results, and analyzing complex data calls for reusable software tools and code (Mitchell2024). There are many data points and low-quality features in metabolomics data, easily leading to artifacts in computational processing (Li2023b). Different from calling nucleotide bases in genomics, metabolomics features are analog signals that require explicit quality metrics. There are clear benefits to separate pre-annotation from annotation. In genomics, a gene sequence can be deposited for many years before annotation is assigned. While the metabolomics community continues working on compound identification, pre-annotation can validate and interpret archived data.

The inflation of compound numbers in database searches can be much reduced by using neutral mass from pre-annotation, not individual m/z features.

The datasets surveyed here return 1∼2,000 high-confidence compounds per method. The number of detected compounds depends on the depth and coverage of analysis. Detection frequency across samples also varies greatly on compounds, raising the question of distinguishing metabolome from exposome (Banbury2025). While the definition of metabolome can be debated, xenobiotics such as medications and environmental pollutants are relevant to human health and should be reported. This study does not address how the coverage of different datasets is affected by experimental methods and conditions, which should be a focus of future computational data processing and annotation. Only human plasma or serum data are analyzed here. Other tissue types may have different complexity, but the approach of “detectable khipu” should be applicable.

## Data and Code Availability

The serial dilution data are on Metabolomics Workbench (https://www.metabolomicsworkbench.org) under the study IDs ST002454. The NIST SRM 1950 data are available at: https://zenodo.org/records/17580076. The 61 asari processed public datasets are available at: https://zenodo.org/records/14541717. The lists of mass patterns for isotopologues, adducts and ISFs are included in the mass2chem software package, freely available at https://github.com/shuzhao-li-lab/mass2chem. All data analysis in this work is provided as Jupyter notebooks at https://github.com/shuzhao-li-lab/dark_metabolome.

## Author Contributions

Project design and writing: SL. Data collection and processing: YC. In-house spectra acquisition: MT, SZ. Data analysis: SL, YC. Software development: JM, YC, SL. All authors participated in and approved the final manuscript.

## Acknowledgements

This work was in part supported by NIH grant R01 AI149746 (NIAID) and ARPA-H award D24AC00345. The content is solely the responsibility of the authors and does not necessarily represent the official views of the funders.

## Supplemental Information

Table S1. List of datasets used in this work, including 45 Orbitrap studies and 16 TOF studies.

Table S2. Most frequent mass differences in Orbitrap ESI+ data.

Table S3. Most frequent mass differences in Orbitrap ESI- data.

Table S4. Most frequent mass differences in TOF ESI+ data.

Table S5. Most frequent mass differences in TOF ESI- data.

Table S6. Isotopologues, adducts and ISFs used for extended khipu search. Table S7. Results of features explained in TOF LC-MS metabolomics data.

File S1. The synthetic dataset by injecting features from authentic compounds into a biological feature table.

## Methods

### Data Retrieval and Processing

A total of 61 public, untargeted metabolomics studies were downloaded from Metabolomics Workbench (metabolomicsworkbench.org) and MetaboLights (www.ebi.ac.uk/metabolights/). All studies were based on human serum or plasma samples, totaling 3482 samples. Of the 61 total studies, 45 were collected using orbitrap-type mass analyzers and 16 using time-of-flight mass analyzers. The breakdown by chromatography and ionization type are noted in Table S1 but all permutations of HILIC, RP, ESI+ and ESI- were represented in all analyzer subsets. Large studies were down selected to between 100∼120 samples.

The raw files were converted to centroided mzML files using ThermoRawFileParser v1.3.1 for Orbitrap data or msConvert v3 for ToF data. Data preprocessing was performed using Asari (v 1.13.1), yielding feature tables with m/z, retention time, peak shape, signal-to-noise ratio, chromatographic selectivity, detection count and intensity values per study. For all analyses, the full feature table was used which enforces the default feature quality filters (signal-to-noise ratio, or SNR, of 2, and a peak shape > 0.5 defined as goodness of fit to a gaussian model). Orbitrap studies were processed using the autoheight option enabled, while ToF data was processed using a mass accuracy of 25 ppm, a min_peak_height of 1000, a cal_min_peak_height as 3e4, and a min_intensity_threshold of 500. Mass calibration was performed according to the default parameters in Asari.

The MoNA MS/MS library were downloaded from https://mona.fiehnlab.ucdavis.edu/ (Feb 8, 2024) and the resulting spectra were deduplicated as follows using utilities from MatchMS (Huber2024). First all spectra for the same precursor inchikey were identified to yield a spectral cluster. Within each spectral cluster, in the second step, the pairwise cosine similarity is calculated for all pairs. The sum of the cosine similarity score weighted by the number of matched peaks for all such comparisons per spectrum is then calculated. Lastly, the spectrum with the largest sum score for that inchikey is then selected as the exemplar for that inchikey and all others are discarded. All collision energy spectra were considered during deduplication with the assumption that the most common fragments should be frequently shared across collision energies.

### LC-MS analysis of glycochenodeoxycholic acid

The chemical standard of glycochenodeoxycholic acid (Cayman Chemical, catalog number 16942) was prepared with acetonitrile at 1 µg/mL. The standard was run using a HILIC column on a Thermo Scientific Q Exactive HF-X Mass Spectrometer (Thermo Fisher Scientific, MA, USA) coupled to Thermo Scientific Transcen LX-2 Duo UHPLC system (Thermo Fisher Scientific, MA, USA), with a HES-II ionization source. The HILIC method utilized an Accucore™ 150 Amide column (10×2.1 mm guard; 100×2.1 mm analytical) with 10 mM ammonium acetate mobile phases (95:5 acetonitrile:water, v/v) containing 0.1% acetic acid, with a gradient from 0% to 98% B2 over 20 minutes at a flow rate of 0.55 mL/min and 45 °C column temperature. MS data were acquired in positive ionization modes (66.7–1000.0 m/z, resolution 60,000 at m/z 200) with optimized ion source parameters, including 3.5 kV spray voltage, 300 °C capillary temperature, and 425 °C heater temperature.

### LC-MS analysis of serial dilution of plasma and SRM 1950 samples

The plasma:H2O serial dilution samples were analyzed on a Thermo Orbitrap ID-X Tribrid mass spectrometer coupled with a HILIC Accucore™ 150 Amide column as previously described in Li et al, (2023b). The data were released as part of the “Bloody Mary” dataset on Metabolomics Workbench. The SRM 1950 dataset was composed from 17 injections from separate batches, using the same above method (Li2023b). The SRM 1950 sample was purchased from NIST (https://www-s.nist.gov/srmors/view_detail.cfm?srm=1950). For blank filtering, a blank sample was taken from each of the 17 separate batches. The resulting samples were processed by asari using the same version and parameters. When a feature in the biological feature table was also detected in the blank and its intensity was no more than 3 times of that in the blank, it was removed from analysis.

### Search of m/z patterns

The isotopic pairs, ^13^C/^12^C features, are defined by annotation of matched retention time, mass distance of 1.003355, and intensity of ^13^C feature under 50% of that of ^12^C feature. The RT differences of all isotopic pairs in each dataset were calculated, and their standard deviation was used to define the RT coelution window per study (one standard deviation on either side of the molecular ion).

To compare MS/MS data to LC-MS data, our deduplicated MoNA MS/MS library is used (13973 precursors in positive and 9184 in negative). A 5-ppm mass tolerance was used for both precursor and MS/MS peak matching with features in retention time window. Only MS/MS peaks whose normalized intensity was higher than 0.1 was considered. To compare common mass difference to LC-MS data, in-source fragment candidates are manually selected from common mass differences, and the maximum of 5 ppm of m/z value or 0.0005 as absolute value was used for mass delta matching. Results here are searched on the top 5 MS/MS fragments, while using top 10 fragments led only to minor increase of matches (not shown).

### Calculation of common mass differences

For the molecular ion in each khipu, the m/z differences to all other features in the elution window were calculated and counted in a histogram (bin size 0.0001 for Orbitrap data, 0.0005 for TOF data). The histograms were smoothed, and peak values were selected by a threshold (100 for Orbitrap data, 20 for TOF data). The top 20 mass differences (deltas) were selected per ionization mode to be used as candidate in-source fragments. Because the histograms use their left edges as reported values, there is a minor shift in some values (e.g. 0.00005 for 1.0033).

The RT shift of these candidate ISFs shows that the majority have identical RT as their molecular ions, confirming coelution in chromatography (**Figure S3b**). The RT shift distributions bear two shoulder peaks, which suggest that a subset of matched ISFs is not truly coeluting, thus not fragments but different compounds. This can lead to an overestimation of ISF contributions.

### Pre-annotation using Khipu

Khipu performs pre-annotation by assigning co-eluting features to adduct and isotopologue relations using a generic tree structure based on *a priori* mass delta patterns (Li2023a). The maximum of 5 ppm of m/z value or 0.0005 as absolute value was used for mass delta matching. The conservative parameters used in Figure 3 without ISFs are as follows: in positive ionization mode, isotope patterns 13C/12C at m/z 1.0034, 13C/12C × 2 at m/z 2.0067, and 37Cl/35Cl at m/z 1.9970; adducts Na/H at m/z 21.9819, ACN at m/z 41.0265, NaCOOH at m/z 67.9874, K/H at m/z 37.9559, and CH3OH at m/z 32.0262. In negative ionization mode, isotope patterns 13C/12C at m/z 1.0034, 13C/12C × 2 at m/z 2.0067, 37Cl/35Cl at m/z 1.9970, and 32S/34S at m/z 1.9958; adducts Na/H at m/z 21.9819, NaCOOH at m/z 67.9874, and NaCH2COOH at m/z 82.0030. The parameters for comprehensive khipu analysis are given in Table S6. The results on TOF datasets are in Table S7.

### Matching compound annotations using neutral mass

The compounds in the genome scale metabolic model (Robinson2020) were retrieved from https://github.com/SysBioChalmers/Human-GEM, and subcellular compartments were removed. The reported metabolites from Mandal et al (2025) were harmonized to consistent HMDB identifiers for this study. Comparisons of khipu neutral masses to HMDB (version 5), Mandal et al (2025) or genome scale metabolic model were based Asari function *asari.tools.match_features*, limited to 5 ppm. Details are given in companion Jupyter notebooks. Without other parameters such as retention time or MS/MS patterns, the matched numbers represent the maximum of probable compound annotation.

### Generating a synthetic feature table using a biological dataset and a compound library

A set of true compounds and their detected features were taken from an authentic compound library, intensity values extracted from the original acquisition. The authentic library was manually verified. The biological data were repeated analysis six SRM1950 samples, using a matched method to the authentic compound library (8-minute HILIC with pos ionization).

The features from authentic compounds were rescaled to match the intensity range in the biological data. As the biological data table contains six samples, each compound is added to a random number of samples, using random intensity redrawn from a gaussian distribution of 50% variance. These synthetic data were injected into the biological data table to become a synthetic data table. If a synthetic feature already existed in the biological data, the biological feature was removed.

## Supplemental Figures

**Figure S1:**
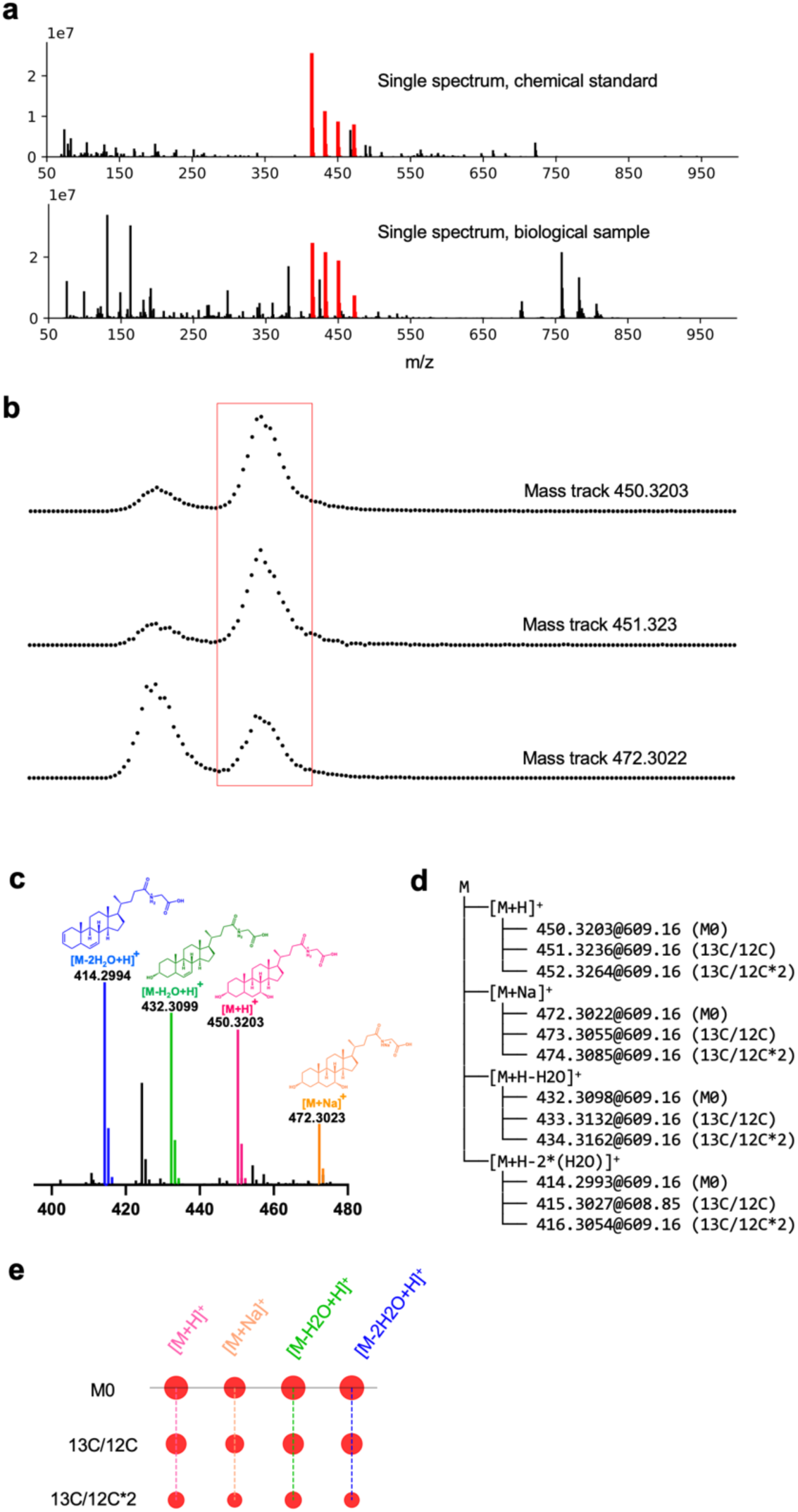
Detailed example of khipu pre-annotation on a bile acid with fragments. This compound has features for two adducts, two neutral losses of H_2_O and three isotopologues. Data are illustrated for both chemical standard and a biological sample. **a.** Full scan mass spectra of glycochenodeoxycholic acid, analyzed from chemical standard (top, containing impurities) and observed in a biological sample (bottom. Sample 214 in study ST002112_RPpos_B3_ppm5_356194, RT 619.49 second). This is chosen to illustrate a complex ion pattern, while most metabolites have fewer ions. **b.** Example mass tracks (extracted ion chromatogram) from same biological sample. Elution profiles are composed from consecutive spectra (scan numbers 600-650), each dot from one spectrum. Red box marks the LC-MS features related to glycochenodeoxycholic acid. Reported retention time may be adjusted by alignment cross samples. **c.** Manual interpretation of the mass peaks (red in **a**). Smaller peaks of the same color are M1 and M2 isotopologues, which are difficult to see in the full mass range in **a**. **d.** Example of khipu pre-annotation of isotopologues, adducts and fragments in a tree structure, with m/z values labeled for each ion. **e.** Khipugram visualization of the data in **d**. Khipu is both the name of software and of a group of ions belonging a putative compound.

**Figure S2:**
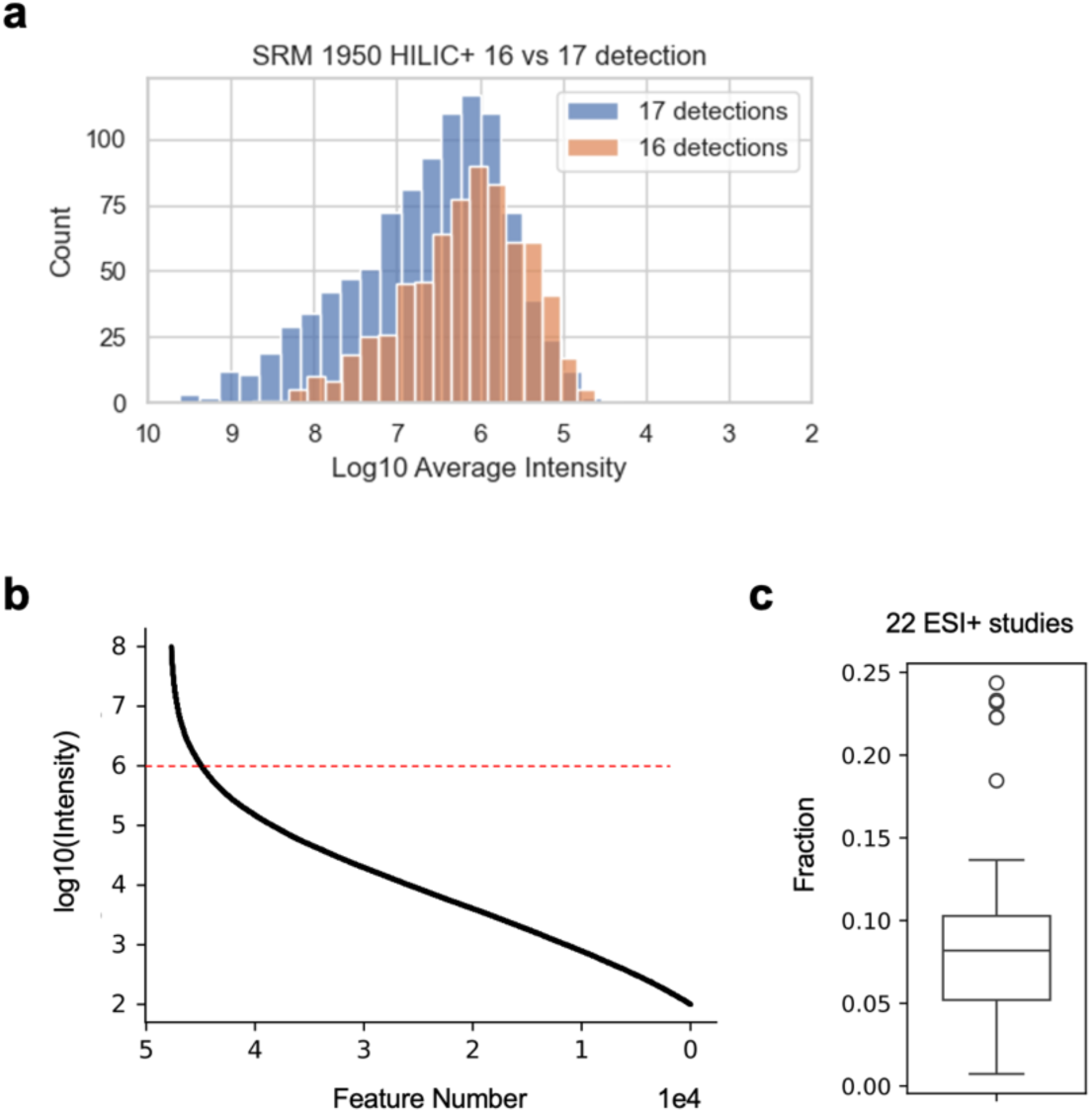
Long-tail distribution of feature intensity in LC-MS metabolomics. **a.** Intensity distributions for the top two feature bins in Figure 1d. **b.** Intensity distribution by feature number, where intensity is plotted at log10 scale. Example from dataset ST002200_RPpos_17min_B3_ppm5_3422144, total 50,647 features. **c.** The fraction of features above 1E6 in intensity cross 22 Orbitrap ESI+ datasets.

**Figure S3:**
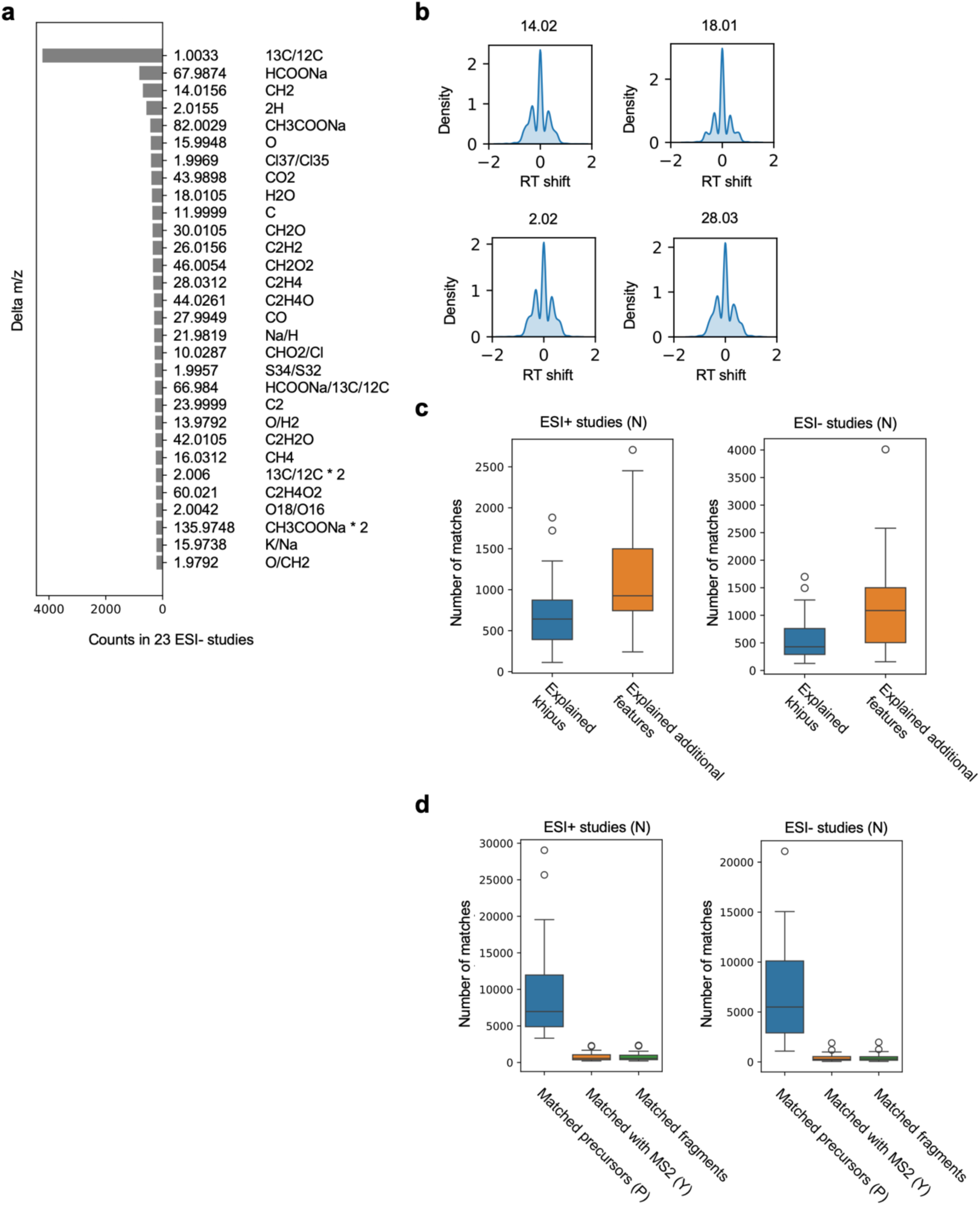
Additional data on common mass deltas and ISFs. **a.** Most frequence delta m/z values in negative ionization Orbitrap datasets. **b.** Example distribution of retention time shift (in seconds) associated with delta m/z (labeled at top). **c.** The numbers of khipus and additional features explained by candidate fragments, in positive and negative ionization datasets, respectively. **d.** Absolute numbers of features matched to MoNA MS/MS spectra across the 45 LC-MS studies, using positive and negative ionization respectively. In each boxplot, 1^st^ column is the number of features matching a precursor m/z value (P), the 2^nd^ column the subset of P with match to at least one MS/MS fragment (Y), and the 3^rd^ column number of all matched MS/MS fragments per study.

**Figure S4:**
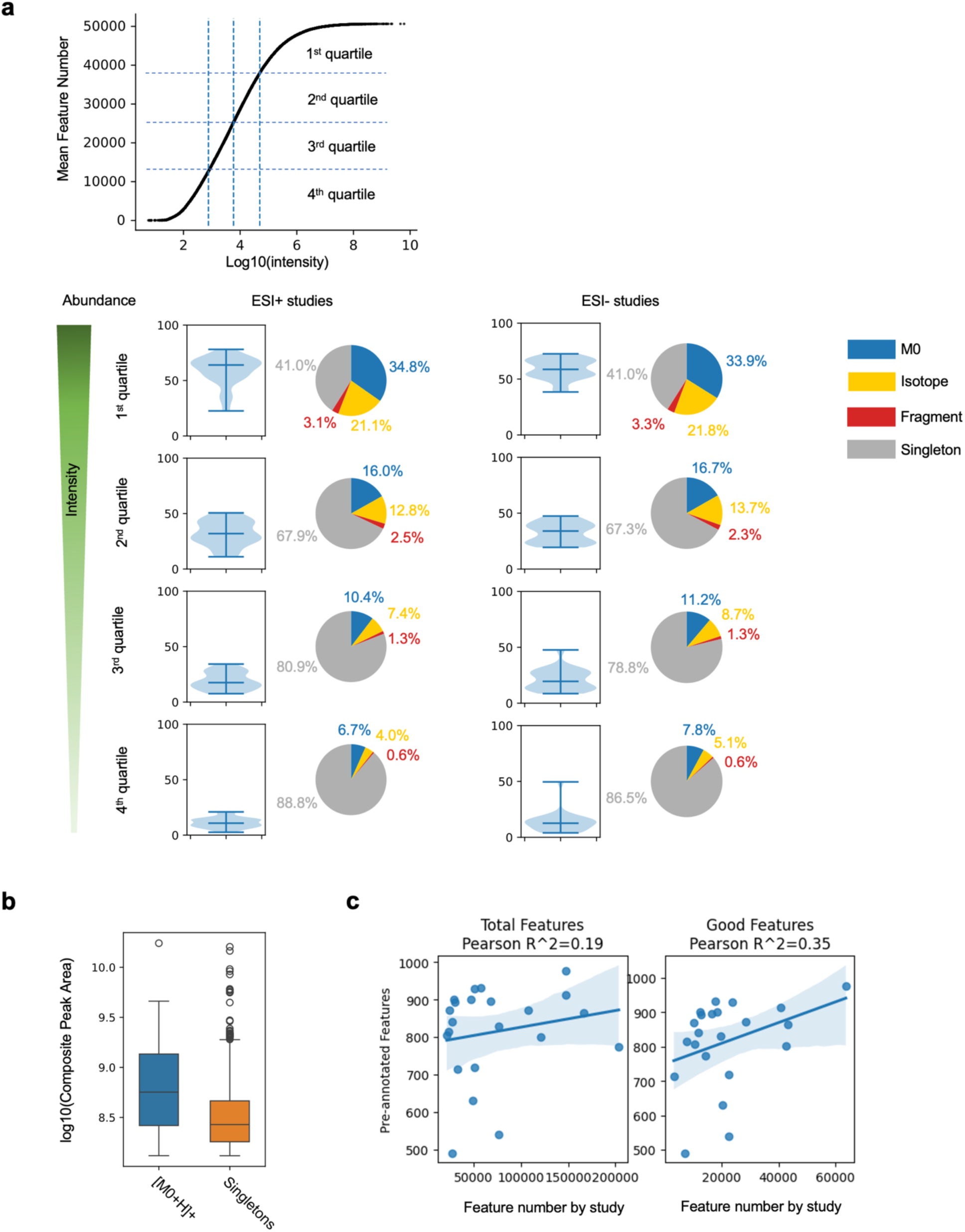
Khipu based pre-annotation explains most of the abundant features. **a.** Top: Example of distribution of feature intensity in Figure S2b divided into quartiles, which are marked by dashed lines. Bottom: Percentage distribution of pre-annotated features in each intensity quartile cross studies, shown as violin plots, for positive and negative ionization datasets respectively. The pie charts show median percentage of contributions from M0, other isotopologues, in-source fragments and singletons. Adducts are included in khipus, and singletons are the unexplained features. **b.** Abundance difference between [M0+H]+ features and singletons within the top quartile. Example from study ST001237_HILICpos_B2_ppm5_3524314, p-value < 1E-25 by Student t-test. **c.** The numbers of khipu explained features in the top bin of Figure 4 (22 Orbitrap ESI+ datasets) in correlation with the total numbers of features (left) and the numbers of good features (SNR > 5 and peak shape > 0.9 when fitting to a gaussian curve; right).

**Figure S5:**
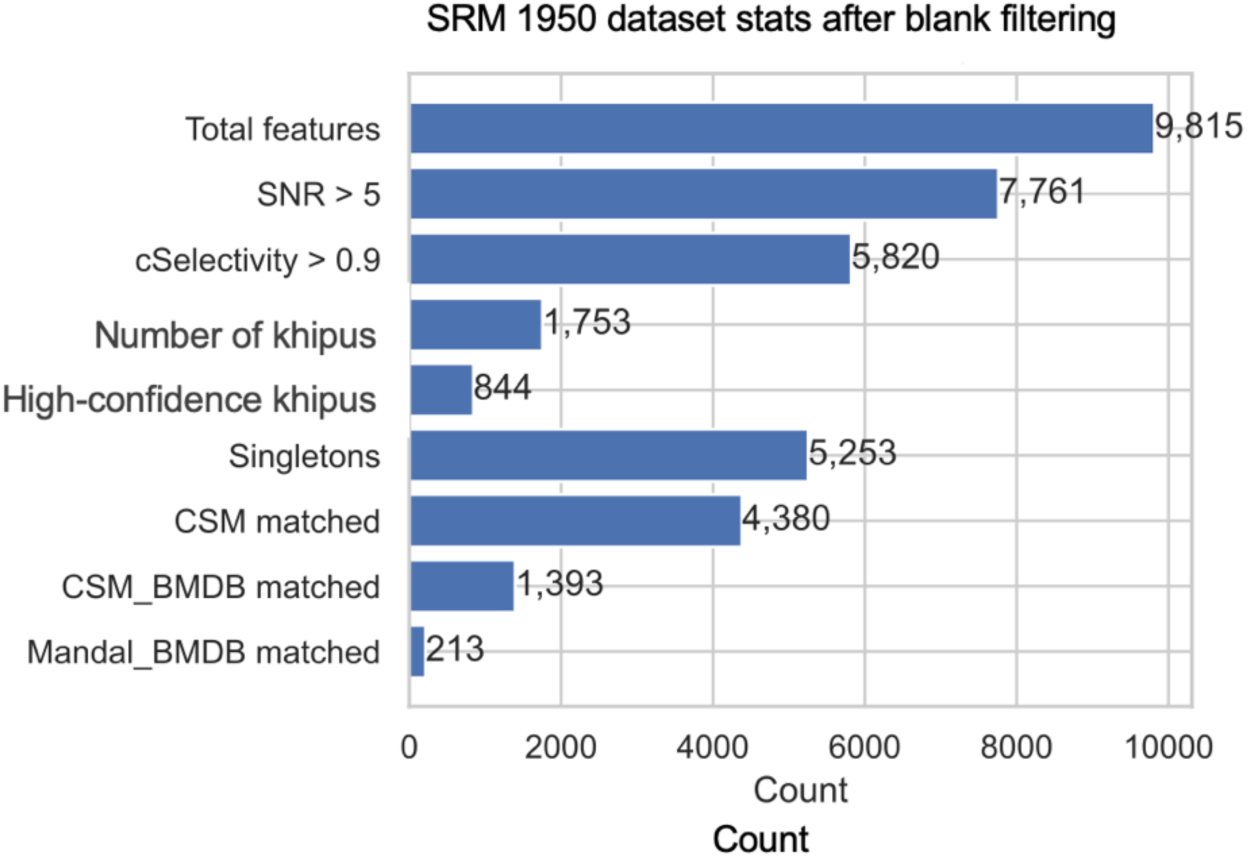
Effect of blank filtering on the annotation coverage. The SRM 1950 dataset in Figure 5c is filtered by features that were measured in companion blank samples in the same experiment.

## Notes

### Competing Interest Statement

The authors have declared no competing interest.

### Summary of Updates

Added new Figure 1, new analyses of dilution series and synthetic data.

